# Polysome-profiling in small tissue samples

**DOI:** 10.1101/104596

**Authors:** Shuo Liang, Hermano Bellato, Julie Lorent, Fernanda Lupinacci, Vincent Van Hoef, Laia Masvidal, Glaucia Hajj, Ola Larsson

## Abstract

Polysome-profiling is commonly used to study genome wide patterns of translational efficiency, i.e. the translatome. The standard approach for collecting efficiently translated polysome-associated RNA results in laborious extraction of RNA from a large volume spread across multiple fractions. This property makes polysome-profiling inconvenient for larger experimental designs or samples with low RNA amounts such as primary cells or frozen tissues. To address this we optimized a non-linear sucrose gradient which reproducibly enriches for mRNAs associated with >3 ribosomes in only one or two fractions, thereby reducing sample handling 5-10 fold. The technique can be applied to cells and frozen tissue samples from biobanks, and generates RNA with a quality reflecting the starting material. When coupled with smart-seq2, a single-cell RNA sequencing technique, translatomes from small tissue samples can be obtained. Translatomes acquired using optimized non-linear gradients are very similar to those obtained when applying linear gradients. Polysome-profiling using optimized nonlinear gradients in HCT-116 cells with or without p53 identified a translatome associated with p53 status under serum starvation. Thus, here we present a polysome-profiling technique applicable to larger studies, primary cells where RNA amount is low and frozen tissue samples.

## INTRODUCTION

Protein levels are modulated via a series of mechanisms including transcription, mRNA-splicing (1), -transport (2), -localization (3), -stability (4), –translation (2) and protein-stability (5). Notably, mRNA translation is the most energy consuming process in the cell (6) and its tight control is therefore essential (7). Consistently, mRNA translation has been suggested to be the predominant post-transcriptional mechanism impacting protein levels (8, 9) although the relative contribution of different mechanisms affecting protein levels is context dependent (10, 11). Moreover dysregulation of translation is associated with diseases including fibrosis (12) and cancer (13). Thus there is a need to study translatomes (i.e. the pool of efficiently translated mRNA) to obtain a more complete understanding of how gene expression is modulated in both health and disease.

Regulation of mRNA translation can be global by affecting mRNAs transcribed from essentially all genes, selective by targeting mRNAs from a gene subset or specific by affecting mRNA copies from a single gene (14). Translation can be divided into four phases: initiation, elongation, termination and recycling (15). Although e.g. elongation is regulated by cellular stress (16) most described modulation of translation occurs at the initiation step where mRNAs are recruited to ribosomes (15, 17). When translational efficiency is regulated via changes in initiation, a change in the proportion of the mRNA copies that are translated (on-off regulation), and/or a continuous shift in the mean number of ribosomes associated with the mRNA copies is observed (18). The mode of regulation appears to, at least partially, depend on the function of the protein encoded by the mRNA as following inhibition of the mammalian/mechanistic target of rapamycin (mTOR), mRNAs harbouring 5’ Terminal Oligopyrimidine Tracts (TOP-mRNAs) in their 5’ un-translated regions (5’UTRs) encoding proteins involved in protein synthesis show on-off regulation while mRNAs e.g. encoding for mitochondria related proteins show continuous shifts in translational efficiency(18). Nevertheless, both modes of regulation leads to alterations in the mean number of ribosomes per mRNA copy and the proportion of the mRNA population that is efficiently translated (e.g. associated with >3 ribosomes)(18). This underlies methods used to assess changes in the translatome.

Polysome-and ribosome-profiling are currently used to study translatomes. Polysome-profiling involves isolation of cytosolic extracts followed by ultracentrifugation on a 5%-50% linear sucrose gradient. During the centrifugation, mRNAs sediment according to how many ribosomes they associate with. Following fractionation, fractions containing efficiently translated mRNAs, e.g. those associated with >3 ribosomes, can be identified and pooled. The RNA-pool is then quantified using either DNA-microarrays or RNA-sequencing (RNAseq) to derive quantitative data on translatomes. The selection of >3 ribosomes to define efficiently translated mRNA is motivated by that most mRNAs appear to involve a change in the amount of such mRNA following a change in their translational efficiency. This is the case for both on-off regulation and continuous shifts due to the nature of the distribution of ribosome association (normal distribution) which leads to that regulation will involve partial transition across the >3 ribosome-association threshold(18). Moreover, the threshold is set such that it captures ∼80% of newly synthesized polypeptides (19, 20). During ribosome-profiling, ribosome protected fragments (RPFs) are generated by applying a mild RNase treatment and isolated using gel purification (21). RPFs are then identified and quantified using RNAseq to reveal nucleotide resolution ribosome positioning. Such data is most commonly used to decipher changes in ribosome location (22, 23), but is also sometimes used to assess changes in translational efficiency(24).

Thus, in contrast to polysome-profiling where efficiently translated mRNAs are quantified (i.e. an mRNA perspective), ribosome-profiling quantifies ribosome-association (i.e. a ribosome-perspective). We recently showed that this leads to a bias when mRNAs showing large changes in translational efficiency (e.g. TOP-mRNAs showing on-off regulation) and those showing smaller continuous shifts (e.g. mitochondrial related mRNAs) are regulated under the same condition. Under these settings ribosome-profiling will to a larger extent as compared to polysome-profiling bias towards identification of abundant mRNAs showing larger changes as differentially translated(18). This is explained by that while a shift of an mRNA from association with 6 ribosomes to 1 ribosome as compared to a shift from 4 ribosomes to 2 ribosomes will be quantified very different when applying ribosome-profiling (3 fold larger difference for a 6-to-1 shift as compared to a 4-to-2 shift), polysome-profiling will assign more similar fold-changes and very similar statistical significance(18). Therefore, for unbiased identification of differential translation under conditions when the precise positioning of ribosomes is not required, polysome-profiling is the preferred technique.

A technical challenge during polysome-profiling is that the pool of efficiently translated mRNA is collected in a large volume (often >3 ml) that is spread across 5-10 fractions (depending on the volume of the gradient and the volume of the collected fractions). This is commonly solved by that mRNA is isolated from each fraction separately using e.g. Trizol followed by pooling during re-suspension of RNA pellets. For small samples, however, such extensive dilution is problematic as it may cause sample loss. Moreover, isolation of mRNA from many fractions is labor intensive and, in larger experimental setups (e.g. large in vitro experimental set-ups or studies of clinical cohorts involving 100s of samples resulting in 1000s of fractions that need to be pooled), may introduce a risk of mistakes such as erroneous pooling of fractions and sample mislabeling. Thus approaches simplifying collection of efficiently translated mRNA (i.e. >3 ribosomes) are warranted. Herein, we describe an optimized non-linear sucrose gradient which collects efficiently translated mRNAs (>3 ribosomes) in only one or two fractions to reduce sample handling 5-10 fold. By coupling isolation of such RNA with quantification methods originally developed for single-cell RNAseq it is possible to derive data on translatomes from small tissue samples such as those collected in biobanks. Importantly, this approach produces very similar data on translatomes as compared to the standard linear gradient approach. Thus, polysome-profiling can now be applied to small samples from tissues or primary cells where RNA amount is often limited.

## MATERIAL AND METHODS

### Preparation of cytosolic lysates from cell lines

HCT116 p53+/+, HCT116 p53-/- and MCF7 cell lines were obtained from ATCC and cultured in DMEM with 10% bovine serum albumin, 1% penicillin/streptomycin and 1% L-glutamine (all from Gibco by Life Technologies). 1 × 10^6^ cells were seeded into a 150×25mm cell culture dish (Corning) two days before harvest to reach a confluency of 70%-80% by the time of the experiment. We assured that expected confluence was reached in each experiment because consistent optimal confluency is critical when assessing translatomes (25). At harvest the media was removed and cell plates were immediately placed on ice followed by an immediate wash with 15 ml ice cold PBS containing cycloheximide (100 μg/ml from Sigma; PBS-CHX). The PBS-CHX was then discarded and cells were detached by scraping, collected in 10ml ice cold PBS-CHX and centrifuged at 266 RCF and 4ºC for 5 min. The cell pellet was re-suspended in 1ml ice cold PBS-CHX, transferred to pre-chilled 1.7 ml tube (Costar) followed by centrifugation at 860 RCF at 4ºC for 3 min. The supernatant was discarded completely and the pellet was re-suspend in 434μl premixed lysis buffer (425μl of 1x hypotonic lysis buffer [50mM Tris-base, 2.5mM MgCl2, 1.5mM KCl in nuclease free water, all from Sigma], 5μl cycloheximide [10mg/ml, Sigma], 1μl DTT [1M, Sigma], 3μl RNaseOUT [Invitrogen by Thermo Fisher Scientific]). Cells were vortexed for 4 sec followed by addition of 25μl of 10% Triton X-100 (Sigma) and 25μl of 10% sodium deoxycholate (Sigma). Lysates were then vortexed for 4 sec and centrifuged at 18727 RCF for 2 mins. 50ul supernatant was collected as cytosolic RNA, diluted with 450ul nuclease free water and 500ul Trizol (Sigma) was added. Cytosolic RNA samples were stored at −80 ºC while lysates were immediately loaded on sucrose gradients.

### Preparation of cytosolic lysates from tissues

All breast cancer samples were from Biobank of A.C Camargo Cancer Center. These were collected under informed consent (ethical permission 1844/13). All tools and tubes were kept in liquid nitrogen to prevent tissue thawing during sample processing. Tissues were pulverized using a BioPulverizer (United Laboratory Plastics) followed by grinding in a liquid nitrogen proof grinding container until the tissue was a fine powder. The tissue powder was collected in a 1.7 ml tube and kept on dry ice. A modified lysis buffer as compared to the one for cultured cells was prepared for tissues where cycloheximide was used at 10-fold higher concentration (i.e. 50 ul of cycloheximide 10mg/ml) and ribonucleoside vanadyl complex [24ul/sample] replaced RNaseOUT. A modified 1000µl tip was used (cut to get a wider entry hole to prevent the tissue powder from clogging) to add 500-1000µl (depending on tissue size) of the ice cold premixed lysis buffer to the tissue followed by mixing by pipetting until the tissue powder homogenously dissolved in the lysis buffer. The sample was transferred to an ice-cold Dounce homogenizer followed by addition of 25μl of 10% Triton X-100 (Sigma) and 25μl of 10% sodium deoxycholate (Sigma); and homogenized on ice by 60 strokes each with the loose and tight pestle. The homogenate was transferred to a chilled 1.7 ml tube and centrifuged at 18727 RCF for 2 min at 4ºC. 50ul supernatant was collected as cytosolic RNA, diluted with 450ul nuclease free water and 500ul Trizol (Sigma) was added. Cytosolic RNA samples were stored at −80 ºC while lysates were immediately loaded on sucrose gradients.

### Preparation of non-linear sucrose gradients

A 5x gradient buffer (20mM HEPES pH 7.6, 100mM KCl 2M, 5mM MgCl2, all from Sigma) was used to prepare 5% (w/v), 34% (w/v) and 55% (w/v) sucrose solutions (1X final concentration gradient buffer (v/v) adjusted with water; all reagents were nuclease free). A gradient cylinder (BioComp) was used to mark each centrifuge tube at the highest level of the cylinder (corresponds to about 5.5 ml when using the Open-Top Polyclear Centrifuge Tubes (14 x 89 mm, SETON Scientific, Part No. 7030). A layering device (BioComp) was used to add 2ml 5% sucrose solution to the bottom of the ultracentrifugation tube; add 34% sucrose solution at the bottom of the tube until the interface between the 5% and 34% solutions reached the marked point; and finally add 55% sucrose solution at the bottom of the tube until the interface between 34% and 55% solutions reached the marked point. The tubes were sealed with rate zonal caps (BioComp) and stored at 4°C for 2 hours before use (to standardize the time between gradient preparation and loading of samples).

### Preparation of linear sucrose gradients

Linear gradients were prepared as described previously(26). Briefly, 5% (w/v) and 50% (w/v) sucrose solutions were prepared in 1x gradient buffer (described above). A gradient cylinder (BioComp) was used to mark the half-full point on each ultracentrifugation tube. A layering device (BioComp) was then used to add 5% sucrose solution until the marked point followed by addition of 50% sucrose from bottom of the tube until the interface between the two solutions reached the marked point. The tubes were sealed with rate zonal caps (BioComp). The gradient maker (BioComp) was then use to generate 5% to 50% linear gradients. Gradients were stored at 4°C for about 2 hours before use (to standardize the time between gradient preparation and loading of samples).

### Sample loading onto sucrose gradients and fractionation

500μl of sucrose solution was removed from the top of the gradient without disturbing gradient composition. About 500μl of cytosolic lysate was then layered on the surface of the gradient. 1x hypotonic lysis buffer was used to balance samples. The samples were centrifuged at 209815 RCF for 2h at 4°C in a TH641 rotor and Beckman Coulter Ultracentrifuge Optima L-90K. Samples were eluted using a gradient station (BioComp). UV was recorded using a BioRad Model EM-1 Econo UV monitor. ∼500ul fractions were collected using a Piston Gradient Fractionator (BioComp) coupled with a fraction collector (Gilson) and the precise location of each fraction along the UV-tracing was monitored using the BioComp software. 750µl of Trizol was continuously added to each collected fraction. Isolated fractions were flash frozen and stored at -80 ºC.

### RNA extraction and purification

Fractions for pooling were identified using the UV tracings recorded by the BioComp software. For linear gradients this corresponded to those fractions with mRNA associated with >3 ribosomes while the fraction containing the center of the peak at the 34% and 55% interface and the peak after were collected from the non-linear gradient (referred to as fraction 0 and +1). These two fractions were pooled and split into two equal parts whereby one was stored and the other was used for RNA extraction. Trizol extraction was performed as described by the manufacturer. Briefly, samples already containing Trizol were thawed on ice. Once thawed, 150μl of chloroform (Sigma) was added to each sample and vortexed for 15 s. The tubes were centrifuged at 18727 RCF for 15 min at 4°C and 750ul of the clear phase was transferred to a new 1.7 ml tube containing 2ul glycoblue (Invitrogen). RNA was precipitated by adding 750ul isopropanol (Sigma), vortexed well and incubated at room temperature for 10 min. Samples were then centrifuged at 18727 RCF at 4°C for 20 min. The supernatant was discarded, the pellet washed with 500ul 70% ethanol and the tube was centrifuged at 18727 RCF for 5 min. The ethanol was removed completely and the pellets were re-suspended in 100μl water (for linear gradients, all pellets for pooling were re-suspended and pooled using the same 100ul water). The resulting RNA samples were cleaned up using the RNeasy MinElute Cleanup Kit (QIAGEN). The RNA quantity was measured by using the Qubit_®_ RNA BR (Broad-Range) Assay Kit and the quality was assessed using the Agilent RNA Nano Chip (Agilent Technologies) on an Agilent 2100 Bioanalyzer.

### Preparation of smart-seq2 RNAseq libraries

Smart-seq2 was performed as described previously using 10ng of RNA as input material (27). RNA sequencing, demultiplexing and conversion to FastQ format was performed by Scilifelab Stockholm. Clustering was done by ‘cBot’ and samples were sequenced on HiSeq2500 (HiSeq Control Software 2.2.58/RTA 1.18.64) with a 1×51 setup using ‘HiSeq SBS Kit v4’ chemistry. Bcl to FastQ conversion was performed using bcl2fastq-1.8.4 from the CASAVA software suite. The quality scale used was Sanger / phred33 / Illumina 1.8+. To ensure that all sequenced data were of sufficient quality, standardized bioinformatics quality control was performed. This included verification of the yield, sequence read quality and cross-sample contamination.

## Analysis of RNAseq data

RNAseq reads were mapped to the human reference genome GRCh38 using Bowtie(28). Two mismatches were allowed in the seed region and reads with multiple possible alignments were excluded. The rpkmforgenes script(29)was used to quantify gene expression (with options-readCount,-fulltranscript and-onlycoding). Genes with zero count(s) were excluded from the analysis. Raw counts were scaled using TMM normalized library sizes(30) and log2 counts per million were computed using the voom function of the limma R package(31). To explore whether the 2 gradient methods led to similar gene expression patterns, principal component analysis was performed after centering by genes. Each sucrose gradient method was then considered separately to assess differential expression in polysome-associated mRNA between HCT-116 p53+/+ and HCT-116 p53−/−cell lines. Differential expression analysis was performed using t-tests applying RVM (Random Variance Model) (32), including the replicate number as factor in the models as implemented in the anota2seq R package. P values were adjusted using the Benjamini-Hochberg (BH) method(33) and a false discovery rate (FDR) < 0.1 was considered significant. The correlation between log2 Fold change between HCT-116 p53+/+ and HCT-116 p53-/- obtained using linear and optimized sucrose gradients was assessed using the Spearman rank correlation coefficient. The p53 translatome identified when using the optimized gradient method was then explored. The optimized gradient polysome-associated mRNA and cytosolic mRNA data were normalized as described above and Anota (34) was used for analysis of differential translation amongst significantly differentially expressed genes on polysome-associated mRNA (using a FDR threshold of 0.2). The replicate number was included as a covariate in the linear models. Unrealistic models of differential translation were excluded (slope>1.5, slope<-0.5) and deltaPT<log2(1.5), deltaP<log2(1.5) were used as filtering criteria of the anota2seqPlotSigGenes function. Genes with an FDR<0.2 were selected for a Gene Ontology (GO) enrichment analysis. Only GO terms with 5 to 500 genes were considered for hypergeometric tests with GOstats (35) where “conditional” was set to FALSE (the structure of the GO graph is not considered in the tests). The 20 top significant GO terms of each set (polysome-associated mRNA up, polysome-associated mRNA down, translation up, translation down) were visualized in a cluster heatmap (row dendrogram shows unsupervised clustering using default method of the gplots: heatmap.2 function(36). All analyses were done using R version 3.3.1.

## RESULTS

### Design of an optimized non-linear sucrose gradient for isolation of efficiently translated mRNA

Because polysome-profiling with linear gradients requires extraction of RNA from many fractions per sample to isolate efficiently translated mRNA (Fig 1A) we considered alternative approaches. As many mRNAs show continuous shifts in translational efficiency within polysomes, pelleting ribosomes would not allow for estimates of their changes in translational efficiencies as such mRNA would be pelleted as long as they are associated with at least 1 ribosome. Instead we reasoned that high percentage sucrose below an intermediate sucrose concentration could potentially enrich for efficiently translated mRNAs (associated with >3 ribosomes) at the surface of the high percentage sucrose – if the optimal sucrose concentrations are identified. This would allow for elution of efficiently translated mRNA in a much smaller volume as compared to the standard linear gradient. While 55% sucrose essentially halts sedimentation we searched to identify a second sucrose concentration that would allow for such enrichment at the 55% sucrose surface. We therefore examined the linear relationship between the log2 number of associated ribosomes and sedimentation distance (which is linearly related to sucrose concentration in a linear gradient)(18). This allowed us to calculate the sucrose concentration which separates mRNAs associated with 3 ribosomes from those associated with 4 ribosomes in a linear gradient (Figure 1B; equals to 34% sucrose). To facilitate entry into the gradient we added a third sucrose layer of 5% sucrose on the top of the 34% sucrose. We next attempted to determine appropriate volumes of each of the sucrose layers. The objective was to position the surface of the 55% cushion close to the top of the tube (to reduce time needed for elution) while still allowing a sufficient volume of 34% sucrose for good separation between efficiently (>3 ribosomes) and less efficiently translated mRNA. A highly reproducible approach for generating layers of sucrose is to start by adding the lowest concentration sucrose to the centrifuge tube and then add increasingly higher sucrose concentrations at the bottom of the tube; thereby pushing lower concentration sucrose solution upwards using the more concentrated sucrose solution(26). While the first layer 5% sucrose can be added by volume directly to the tube, additional volumes for layers are best determined by monitoring the interface between layers and let these reach a certain position in the tube. It is essential that this position can be reproducibly indicated on the tube and we therefore used the same approach as when making a linear gradient using the BioComp gradient maker whereby a cylinder is used to indicate the desired level on the tube(26). For the centrifuge tubes we used, this corresponds to about 5.5 ml. Our initial test of the optimized non-linear gradient indicated separation of 40S, 60S ribosomal subunits and the 80S monosome followed by a peak at the interface between the 34% and 55% sucrose (Fig 2A).

**Figure 1.**
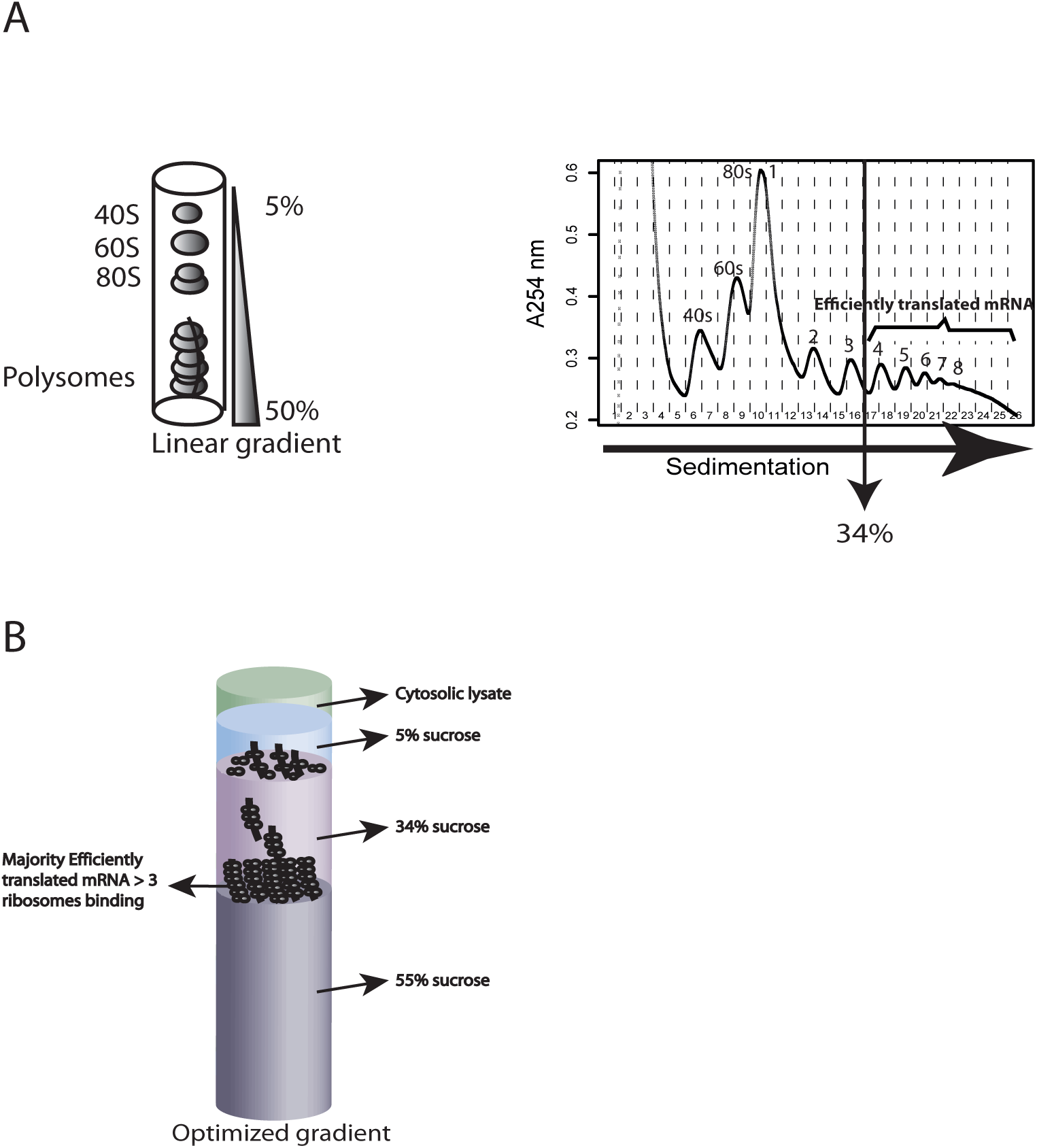
**A**. Polysome profiling using a linear 5% to 50% sucrose gradient. Cytoplasmic RNA isextracted and loaded on the linear gradient. Following ultracentrifugation 40S and 60S ribosome subunits; the 80S monsomes; and polysomes are separated (schematics show UV tracing at 254 nm across the sucrose gradient). Efficiently translated mRNA are isolated from fractions containing mRNA associated with more than 3 ribosomes. This corresponds to a sucrose concentration of 34%. **B**. The optimized non-linear sucrose gradient is made of layers of 5% sucrose, 34% sucrose and 55%sucrose. The purpose is to collect mRNA associated with more than 3 ribosomes at the 55% sucrose surface.

**Figure 2.**
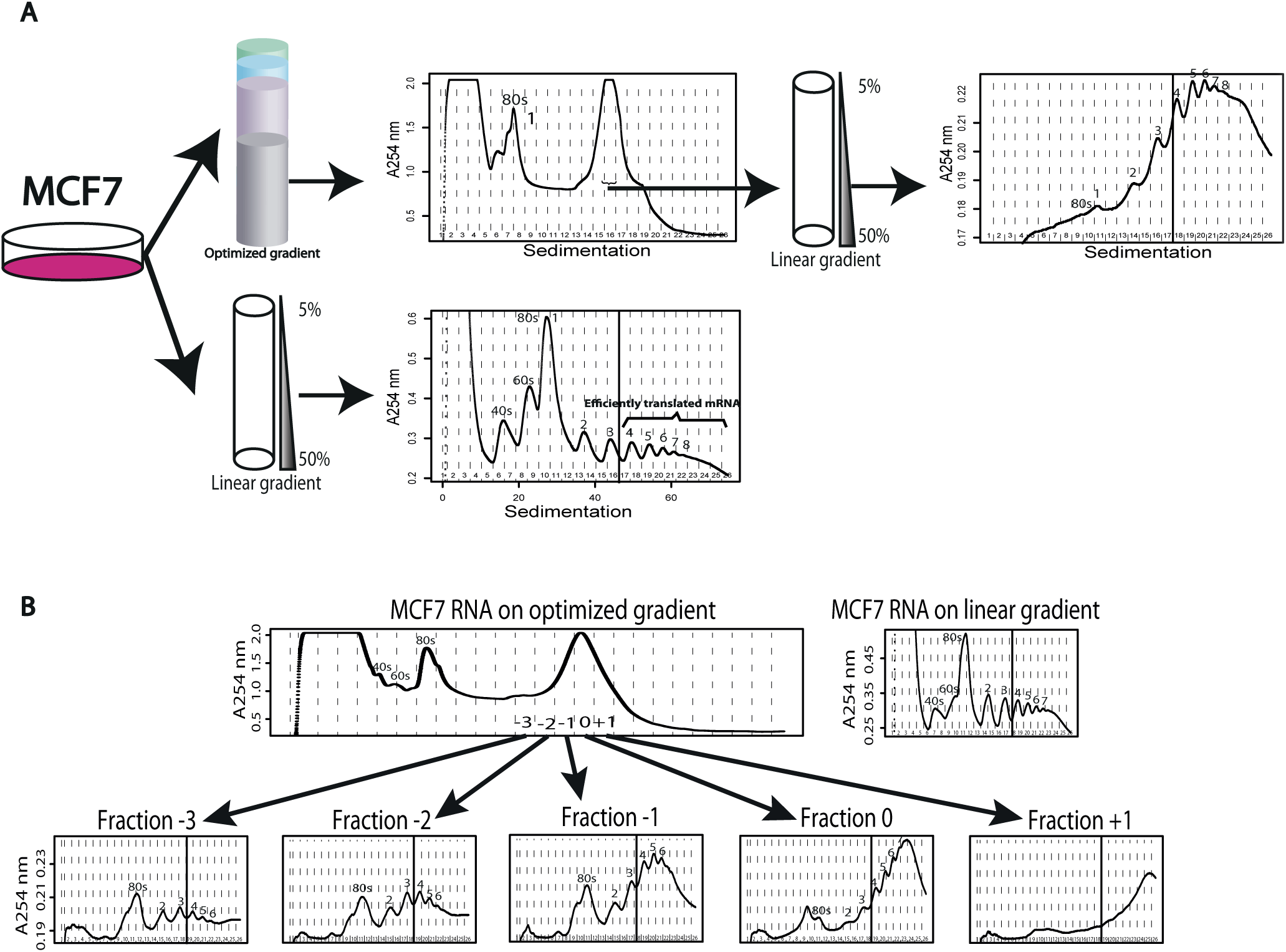
Identification of non-linear sucrose gradient fractions that reproducibly contain mRNAassociated with >3 ribosomes. **A**. Cytosolic lysate from MCF7 cells was sedimented on the optimized non-linear sucrose gradient or a linear sucrose gradient in parallel. The nature of the large high peak was explored using a linear sucrose gradient. This indicated a strong enrichment of mRNA associated with >3 ribosomes as compared to what is expected from running the extract on the linear gradient. **B**. The same approach was used but multiple fractions from the optimized non-linear gradient were evaluated using linear gradients. The fraction containing the center of the peak was designated 0 while fractions towards the bottom or top of the gradient were assigned positive or negative numbers respectively. Shown is also the profile from a linear gradient processed in parallel.

### Reproducible isolation of efficiently translated mRNA using the optimized gradient

We noticed that the width of the peak at the interface between the 34% and 55% sucrose layers varied with the time between preparation of the gradient and centrifugation of the sample. We therefore standardized the time between these to 2 h (which is more than sufficient to prepare lysates to load onto the gradient). This suggests that a small gradient is formed at the interface between the 34% and the 55% solutions which can be standardizing by fixing the time lag. We next sought to explore the nature of the fractions surrounding the large peak to assess whether this peak indeed is enriched for mRNA associated with >3 ribosomes. To this end we used a large batch of cells (12 plates [15 cm] of MCF7 cells) and sedimented their lysate on the optimized non-linear gradient. The major area of the peak between the 34% and the 55% sucrose was spread across 3 fractions. We designated the fraction containing the center of the peak as 0 and those towards the 34% sucrose −1 (etc.) while those towards the 55% sucrose +1 (etc.). The vast amount of RNA used as input allowed us to collect 5 fractions surrounding the peak between the 34% and the 55% sucrose solutions and dilute these 7-fold to enable a run of each fraction separately on the standard 5%-50% linear sucrose gradient (these samples would sink into the linear gradient without dilution). In addition we generated a new cytosolic lysate (from 2 additional plates of MCF7 cells which were cultivated in parallel with those used for the non-linear gradient) and loaded this on a linear gradient as control. As shown in Figure 2B, fraction 0 and fraction +1 are strongly enriched for mRNA with >3 ribosomes while fractions <0, while still being enriched for efficiently translated mRNA, show relatively more mRNA associated with fewer ribosomes. This pattern was obtained in two additional independent experiments (Figure S1). We therefore concluded that collection of the fractions containing the center of the peak and the one further towards the 55% sucrose allows for isolation of efficiently translated mRNA (>3 ribosomes). Thus efficiently translated RNA can be obtained by collecting one fraction (by using the +1 fraction) or two fractions (by using the 0 and the +1 fractions). We deployed a strategy where two fractions (i.e. 0 and +1) are pooled and then split into two tubes whereby one tube is used for RNA isolation while a second tube is stored as a backup. This reduces that risk of losing samples due to failed RNA isolation. Moreover, it allows for downstream processing of only one tube containing efficiently translated mRNA.

### The optimized non-linear gradient allows for consistent isolation of high quality RNA

A concern is that application of the optimized non-linear gradient leads to a reduction in the amount of isolated efficiently translated mRNA. To assess this, and to validate that high quality RNA can be reproducibly obtained, we used two cells lines that differ in their p53 status (HCT116p53+/+ and HCT116p53-/-). We applied serum starvation to these cells (16 h) as translation is commonly modulated during cellular stress and this setup would allow us to assess whether p53 affected such responses (see below). Serum starvation had a comparable effect on global translation for the two cell lines as judged by the similar reductions in polysome-associated RNA coupled with an increase in sub-polysomal RNA (Fig. S2). To allow for a rigid comparison between the standard linear gradient and the optimized non-linear gradient we prepared cytosolic lysates from 6 plates from each cell type and divided the lysates equally between the optimized non-linear gradient and the linear gradient We then collected fractions corresponding to RNA associated with >3 ribosomes from the linear gradient and peak 0 and +1 (as defined above) from the optimized non-linear gradient. We repeated the experiment four times and measured the RNA quantity. As shown in figure 3A, the optimized nonlinear gradient and linear gradient allowed for isolation of similar amounts of efficiently translated mRNA. We then used the Agilent Bioanalyzer to assess RNA quality quantified by RNA Integrity Numbers (RINs; the scale range from 10 [perfectly intact RNA] to 1 [completely degraded RNA]). This revealed that both gradients allow for consistent isolation of essentially perfectly intact RNA (Figure 3B). Thus the optimized non-linear gradient shows similar performance as compared to the linear gradient.

**Figure 3.**
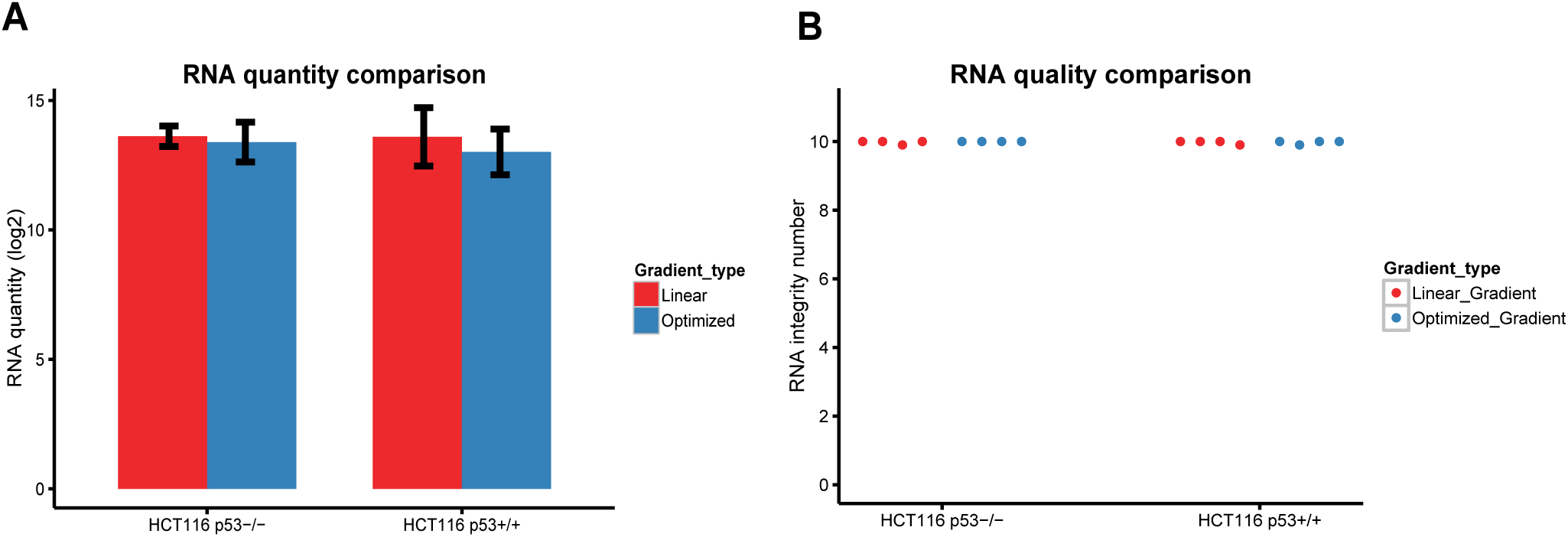
Comparison quantity and quality of RNA isolated using the optimized non-linear sucrosegradient or the linear sucrose gradient. **A**. Mean of total amount of RNA (log2 ng) from quadruplicate experiments are shown in red (Linear gradient) and blue (Optimized gradient) for both HCT116 p53+/+ and HCT116 p53-/- cells. **B**. A scatter plot indicating the RNA integrity number (RIN) from the quadruplicate experiments. Red dots represented linear gradient and blue dots were optimized gradient for both HCT116 p53+/+ and HCT116 p53-/- cell lines.

### The optimized non-linear gradient or the linear gradient generate similar translatomes

A key property of a method that strives towards studies of translatomes using isolation of efficiently translated mRNA (>3 ribosomes) is that the generated data are similar to those obtained using the standard approach (employing linear gradients). To evaluate this, we determined translatomes from serum starved (16 h) HCT-116 cells with and without p53 (as described above). To mimic a situation observed in tissue samples or primary cells where obtained RNA amounts are often small, we employed the smart-seq2 protocol for generation of RNAseq libraries that was developed for single-cell RNA-seq. Notably, this protocol allows for generation of libraries with similar coverage and robustness as standard approaches using <10 ng of input RNA but can generate libraries using <1ng of starting total RNA(27). We therefore generated smart-seq2 libraries using 10 ng of RNA as input from efficiently translated mRNA obtained using standard gradients; optimized non-linear gradients; and cytosolic RNA (i.e. input RNA that was loaded on the gradients). Following normalization, we used principal component analysis to explore the data set. As expected in analysis of polysome profiling data, the first component containing the main source of variation originates from RNA source such that cytosolic mRNA samples separate from polysome-associated mRNA samples. The second and third principal components group samples according to replicate number and p53 status respectively (Fig. 4A). This is consistent with that the gradients produce similar data on translatomes. To further substantiate this conclusion we compared gene expression between HCT-116 p53+/+ and p53−/− cells separately using polysome-associated mRNA isolated from the optimized sucrose gradient or polysome-associated mRNA isolated from a standard linear gradient. At an FDR threshold of 0.1, the optimized gradient approach identified more differentially expressed genes as compared to the linear gradient method (Fig. 4B) but the overlap between the two methods was almost complete (Fig. 4C). Furthermore, the fold-changes obtained between HCT-116 p53+/+ and p53−/− cells when applying the two approaches showed high correlation (Spearman coefficient: 0.75, Fig. 4D). Strikingly, the differences between the two techniques was the obtained FDRs rather than in fold changes such that lower FDRs were obtained when using the optimized gradient (Fig. 4E). We therefore conclude that the optimized non-linear gradient can be applied to obtain translatomes which are similar to those obtained using standard linear gradients. Furthermore, data generated from the optimized sucrose gradient method allowed us to identify the p53 translatome of HCT-116 cells. This revealed that p53 status affects both mRNA levels and translational efficiencies (Fig. 5A). Moreover, genes regulated by these two modes were enriched for distinct cellular functions which is consistent with that p53 reprograms cell phenotypes via modulation transcription and translation.

**Figure 4.**
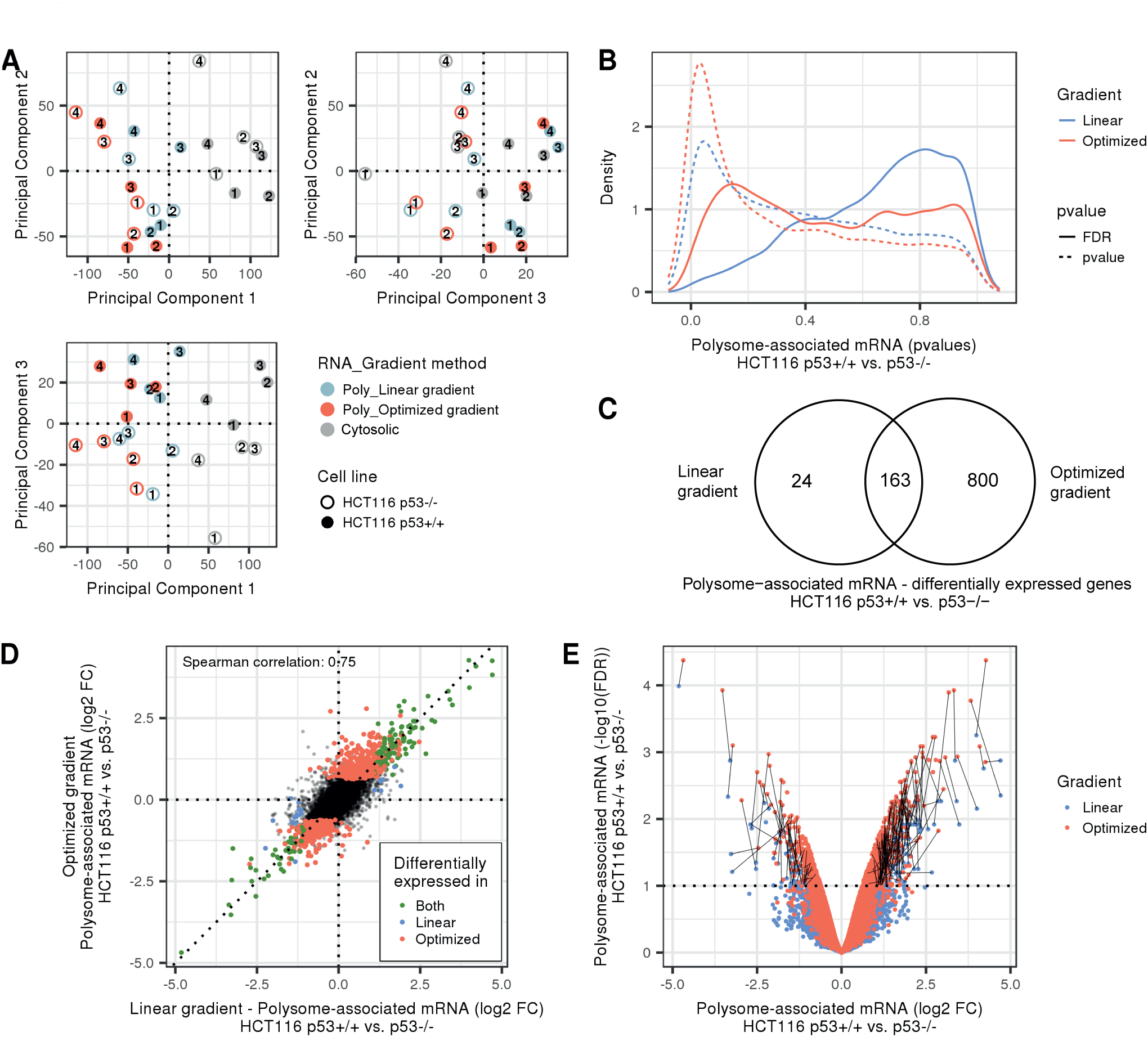
Comparison of linear and optimized sucrose gradient for analysis translatomes. **A**. Projection of all samples in the three first components of the principal component analysis. Replicate numbers are indicated in black. **B-E**. For both gradient methods, an analysis of differential expression on polysome-associated mRNA between p53+/+ HCT-116 and p53-/- HCT-116 cell lines was performed. **B**. Density plot of pvalues (dashed lines) and false discovery rate (plain lines) for both gradient methods (linear gradient in blue, optimized gradient in red). **C**. Venn diagram showing the overlap of genes identified using both gradient methods (mRNAs with an FDR<0.1 were considered differentially associated with polysomes). **D**. Scatter plot showing log2 fold changes using data from optimized non-linear gradient vs. the linear gradient. Different colours correspond to genes identified as differentially polysom-associated by both gradient methods (green); the linear gradient only (blue); the optimized gradient only (red); or none of the methods (black). E. Volcano plot (-log10 FDR vs. log_2_ Fold change) for each gradient method (linear gradient in blue, optimized gradient in red). For genes considered significant by both methods, a black lines join the results per mRNA. Poly = Polysome-associated mRNA; FDR = False Discovery Rate; FC = Fold Change

**Figure 5.**
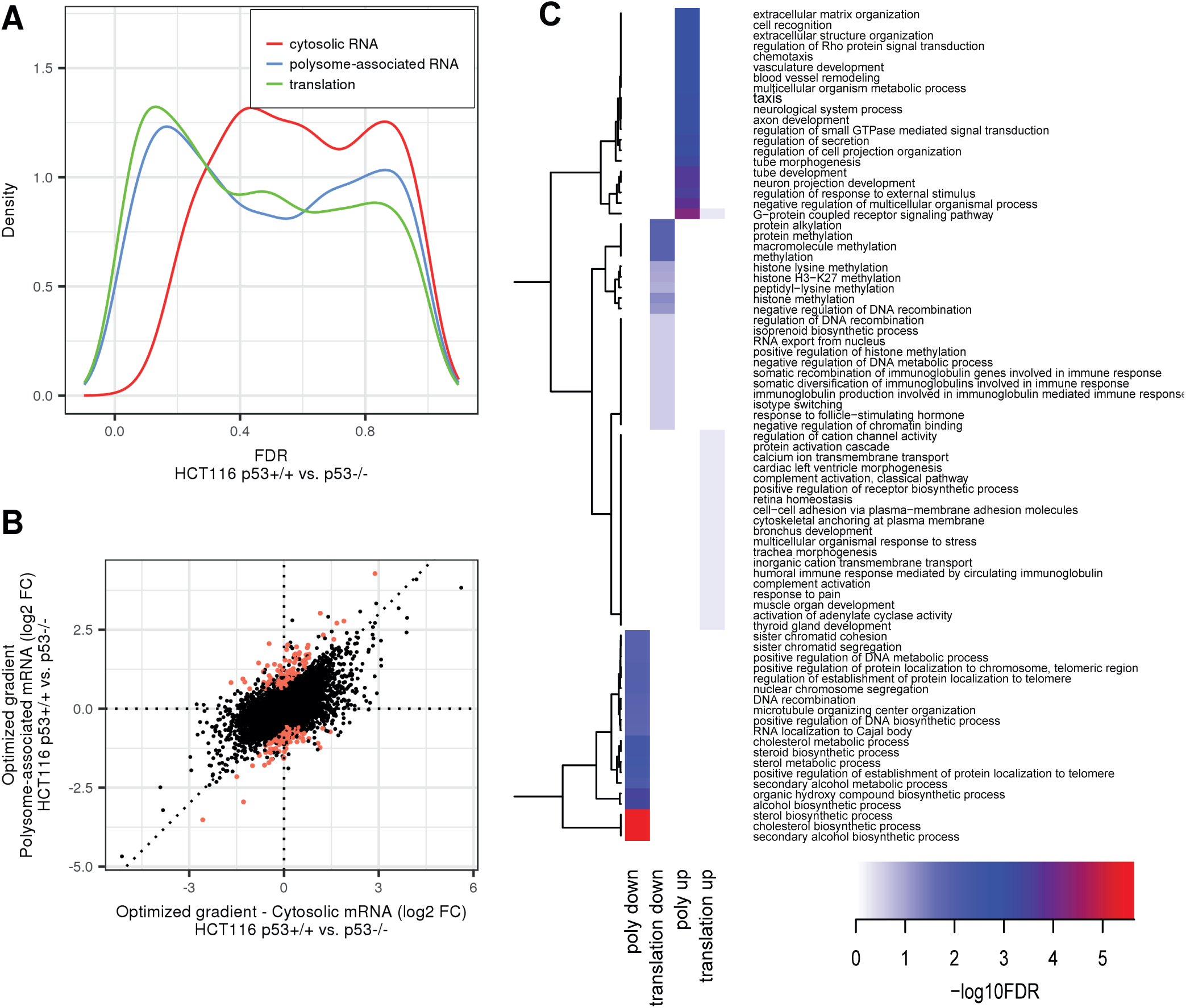
Analysis of the p53 translatome of HCT-116 cells using an optimized gradient. **A**. Density plot of FDR for analysis of polysome-associated mRNA, cytosolic mRNA and analysis of translation. B. Scatter plot of polysome-associated mRNA log2 fold changes of vs. cytosolic mRNA log2 fold changes. Genes significantly translationally regulated are indicated in red. C. Cluster heatmap showing GO term enrichment in genes regulated via changes in polysome-associated mRNA (poly up or poly down) or genes which are translationally regulated(translation up or translation down). –log10 FDR of the hypergeometric tests are colour-coded in the heatmap and an unsupervised clustering was applied to GO genesets. FDR = False Discovery Rate; FC = Fold Change.

### Polysome-profiling of bio-banked breast cancer tissues using optimized non-linear gradients and smartSeq2

To comprehensively evaluate the non-linear sucrose gradient for isolation of efficiently translated mRNA from bio-banked tissue samples we identified a cohort of 161 breast cancer tissues and applied the optimized non-linear gradient to isolate their efficiently translated mRNA. The sizes of the tissue samples were not recorded in the bio-bank but, as estimated by eye, varied between <30 to ∼100 mg. The profiles obtained from the non-linear gradient varied (Fig. S3), likely affected by the RNA integrity in the tissues, but we could nevertheless consistently identify the 0 and +1 fractions which were pooled and split to generate one sample for RNA isolation and one that was stored as backup. We then assessed the quality of the isolated RNA in cytosolic extracts (i.e. input) and following isolation of efficiently translated mRNA using optimized non-linear gradients. This indicated a strong relationship between RINs obtained in input RNA and those obtained for efficiently translated mRNA (Fig 6A). RINs for efficiently translated RNA were larger (i.e. indicating higher RNA integrity) as compared to those observed in input samples (Fig 6B). This shows that a low RIN for the pool of efficiently translated RNA is not caused by the isolation technique, but rather by lower initial RNA quality in those tissue samples. It also highlights that polysome-profiling enriches for intact mRNA. Many of these samples generated very low RNA amounts (<1ng/ul and <10 ng in total) and hence application of protocols adopted for single cell sequencing was essential to generate translatomes. Application of Smart-seq2 to a subset of these samples generated RNAseq libraries amendable for RNAseq (Fig S3). Thus, combining the optimized non-linear gradient with smart-seq2 allows for exploration of translatomes in small tissue samples from bio-banks.

**Figure 6.**
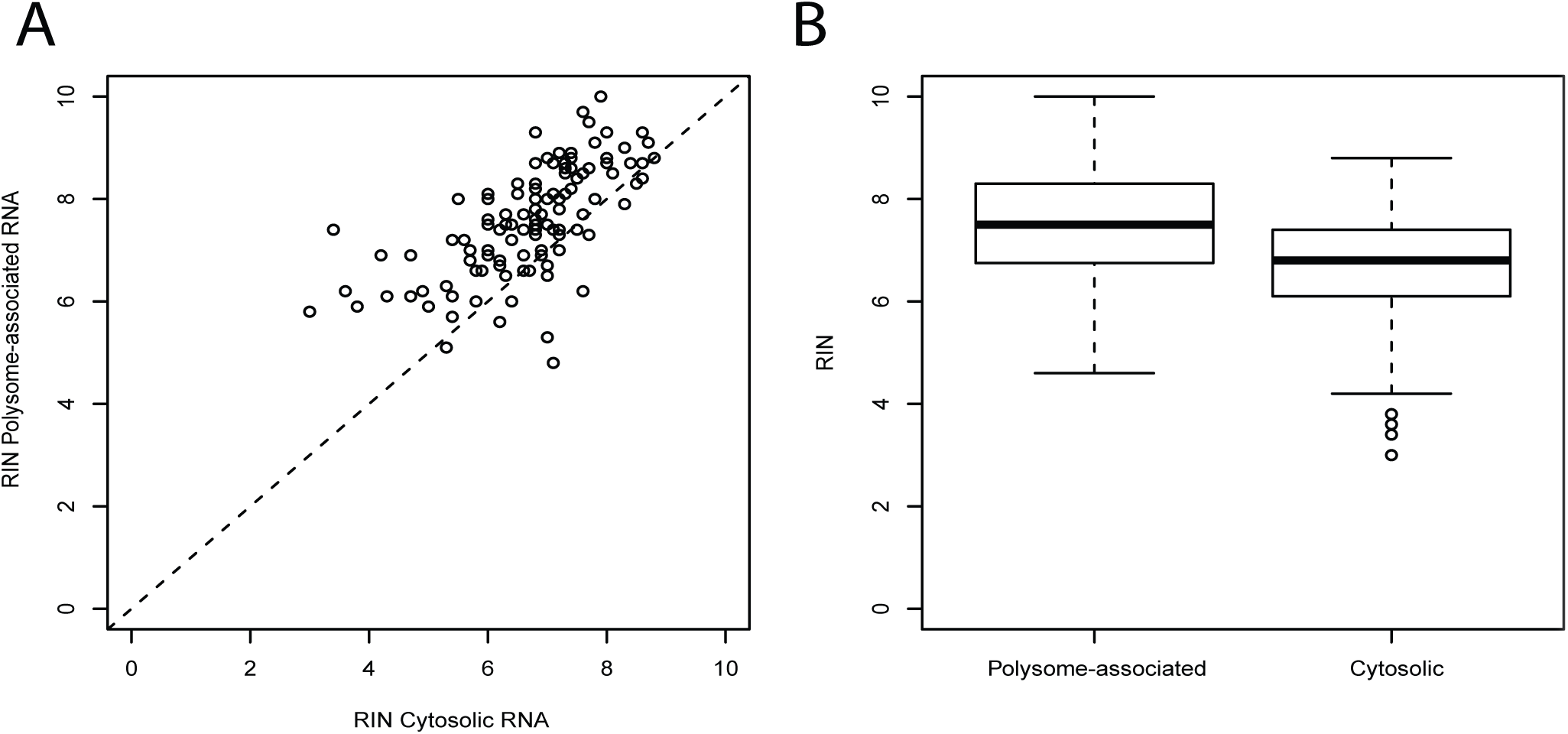
Polysome-profiling of bio-banked breast cancer tissues using optimized non-linear gradients and smartSeq2. A. Scatter plot of RNA integrity number (RIN) obtained from polysome-associated mRNA samples vs. cytosolic mRNA samples when using the optimized. B. Boxplot of RIN for polysome-associated and cytosolic mRNA of the same samples as in A.

## DISCUSSION

Many studies show that translational control can have dramatic effects on the proteome without changes in mRNA levels (17). This was for example observed following various stresses and following modulation of the activity of key cellular pathways such as the mTOR pathway (14). Given the pivotal role of cell signaling and cell stress during human diseases, the translatome is expected to be modulated under a range of pathological conditions. Yet, translatomes are vastly understudied as compared to transcriptomes (14). This reflects that while genome wide approaches for measurements of transcriptomes are relatively easy to apply, methods for studying translatomes have not scaled well to larger sample collections. Although polysome-profiling is the preferred method for studies of changes in translational efficiency as compared to ribosome-profiling when the precise location of ribosomes is of lesser importance (18) (due to the bias introduced when applying ribosome-profiling as discussed above), both these techniques are highly laborious and therefore challenging to apply in large studies. Here we focused on one aspect of polysome-profiling which makes it inconvenient namely that the pool of efficiently translated RNA is obtained in a large volume distributed across multiple fractions that need to be collected and pooled (26). When studying translatomes in small tissue samples or primary cells (37, 38) this poses an additional limitation by leading to extensive dilution of the efficiently translated mRNA which may cause sample loss. We addressed these issues by introducing an optimized non-linear sucrose gradient which allows for consistent isolation of efficiently translated mRNA associated with >3 ribosomes in only one or two fractions. The ability to isolate efficiently translated mRNA enriched for mRNA associated with >3 ribosomes is important to capture changes in translational efficiency manifested by continuous shifts in translational efficiency within polysomes. Changes in translation of such genes will be more difficult to detect using e.g. pull-down approaches as association with one ribosome is sufficient for pull-down of the RNA. As a result such pull-down approaches will likely be biased towards identification of mRNAs showing on-off regulation.

Here we introduced single-cell sequencing protocols for quantification of efficiently translated mRNA. Similar to when obtaining translatomes from limited numbers of primary cells (38), this is essential to allow for studies of translatomes using bio-bank tissue samples as the amount of RNA isolated from such samples is insufficient for standard RNAseq protocols.

Although the presented approach alleviates some of the issues with polysome-profiling it does not circumvent the need of sucrose gradients which is still a limiting factor when planning large studies of translatomes. Nevertheless, this approach made it possible for us to process >150 clinical samples. Thus, polysome-profiling can now be applied to large collections of small tissue samples or primary cells.

## ACKNOWLEDGEMENTS

We acknowledge support from Science for Life Laboratory, the Knut and Alice Wallenberg Foundation, the National Genomics Infrastructure funded by the Swedish Research Council, and Uppsala Multidisciplinary Center for Advanced Computational Science for assistance with massively parallel sequencing and access to the UPPMAX computational infrastructure.

## FUNDING

This research was supported by the Swedish Research Council, the Swedish Childhood Cancer Foundation, the Swedish Cancer Society, the Cancer Society in Stockholm, the Wallenberg Academy Fellows Program, and STRATCAN grants (O. L.). L. M. is supported by a post-doctoral fellowship from the Swedish Childhood Cancer Foundation.

---

**Supplementary figure 1.**. Identification of non-linear sucrose gradient fractions that reproducibly contain mRNA associated with >3 ribosomes. Two additional replicate experiments as performed in Fig. 2B.

**Supplementary figure 2. A.**. Western blot of p53 expression in HCT116 p53+/+ and HCT116 p53-/- cell with or without serum starvation. **B**. Effects of serum starvation on global translation as assessed by polysome-tracings using linear gradients. Shown are overlays for HCT116 p53+/+ and HCT116 p53-/- with or without serum. Amount of efficiently translated mRNA was quantified from peak 4 on (from mRNA bounded with 4 complete ribosomes to more) to the end of the profile.

**Supplementary figure 3.**. Polysome-profiling of tissue samples using the optimized non-linear gradient and smartSeq2. Shown are five examples of polysome profiles from breast cancer tissue samples sedimented on the optimized non-linear sucrose gradient together with their corresponding smartSeq2 RNA sequencing libraries (both cytosolic input and RNA isolated from fractions 0 and +1). For each tumour the polysome-tracing and bioanalyzer tracings showing the size distribution of the smartSeq2 libraries following tagmentation are indicated.

## REFERENCES

1 Bava, F.A., Eliscovich, C., Ferreira, P.G., Minana, B., Ben-Dov, C., Guigo, R., Valcarcel, J. and Mendez, R. (2013) CPEB1 coordinates alternative 3’-UTR formation with translational regulation. Nature, 495, 121–125.

2 Rousseau, D., Kaspar, R., Rosenwald, I., Gehrke, L. and Sonenberg, N. (1996) Translation initiation of ornithine decarboxylase and nucleocytoplasmic transport of cyclin D1 mRNA are increased in cells overexpressing eukaryotic initiation factor 4E. Proc Natl Acad Sci U S A, 93, 1065–1070.

3 Holt, C.E. and Schuman, E.M. (2013) The central dogma decentralized: new perspectives on RNA function and local translation in neurons. Neuron, 80, 648–657.

4 Lindstein, T., June, C.H., Ledbetter, J.A., Stella, G. and Thompson, C.B. (1989) Regulation of lymphokine messenger RNA stability by a surface-mediated T cell activation pathway. Science, 244, 339–343.

5 Zackrisson, S., Janzon L Fau − Manjer, J., Manjer J Fau − Andersson, I. and Andersson, I. Improved survival rate for women with interval breast cancer-results from the breast cancer screening programme in Malmo, Sweden 1976–1999.

6 Roux, P.P. and Topisirovic, I. (2012) Regulation of mRNA translation by signaling pathways. Cold Spring Harb Perspect Biol, 4.

7 Morita, M., Gravel, S.P., Hulea, L., Larsson, O., Pollak, M., St-Pierre, J. and Topisirovic, I. (2015) mTOR coordinates protein synthesis, mitochondrial activity and proliferation. Cell Cycle, 14, 473–480.

8 Kristensen, A.R., Gsponer, J. and Foster, L.J. (2013) Protein synthesis rate is the predominant regulator of protein expression during differentiation. Mol Syst Biol, 9, 689.

9 Schwanhausser, B., Busse, D., Li, N., Dittmar, G., Schuchhardt, J., Wolf, J., Chen, W. and Selbach, M. (2011) Global quantification of mammalian gene expression control. Nature, 473, 337–342.

10 Jovanovic, M., Rooney, M.S., Mertins, P., Przybylski, D., Chevrier, N., Satija, R., Rodriguez, E.H., Fields, A.P., Schwartz, S., Raychowdhury, R. et al. (2015) Immunogenetics. Dynamic profiling of the protein life cycle in response to pathogens. Science, 347, 1259038.

11 Liu, Y., Beyer, A. and Aebersold, R. (2016) On the Dependency of Cellular Protein Levels on mRNA Abundance. Cell, 165, 535–550.

12 Parker, M.W., Rossi, D., Peterson, M., Smith, K., Sikstrom, K., White, E.S., Connett, J.E., Henke, C.A., Larsson, O. and Bitterman, P.B. (2014) Fibrotic extracellular matrix activates a profibrotic positive feedback loop. The Journal of clinical investigation, 124, 1622–1635.

13 Bhat, M., Robichaud, N., Hulea, L., Sonenberg, N., Pelletier, J. and Topisirovic, I. (2015) Targeting the translation machinery in cancer. Nat Rev Drug Discov, 14, 261–278.

14 Piccirillo, C.A., Bjur, E., Topisirovic, I., Sonenberg, N. and Larsson, O. (2014) Translational control of immune responses: from transcripts to translatomes. Nat Immunol, 15, 503–511.

15 Hershey, J.W., Sonenberg, N. and Mathews, M.B. (2012) Principles of translational control: an overview. Cold Spring Harb Perspect Biol, 4.

16 Leprivier, G., Remke, M., Rotblat, B., Dubuc, A., Mateo, A.R., Kool, M., Agnihotri, S., El-Naggar, A., Yu, B., Somasekharan, S.P. et al. (2013) The eEF2 kinase confers resistance to nutrient deprivation by blocking translation elongation. Cell, 153, 1064–1079.

17 Ruggero, D. (2013) Translational control in cancer etiology. Cold Spring Harb Perspect Biol, 5.

18 Gandin, V., Masvidal, L., Hulea, L., Gravel, S.P., Cargnello, M., McLaughlan, S., Cai, Y., Balanathan, P., Morita, M., Rajakumar, A. et al. (2016) nanoCAGE reveals 5’ UTR features that define specific modes of translation of functionally related MTOR-sensitive mRNAs.Genome Res, 26, 636-648.

19 Warner, J.R., Knopf, P.M. and Rich, A. (1963) A multiple ribosomal structure in protein synthesis. Proceedings of the National Academy of Sciences of the United States of America, 49, 122–129.

20 Gierer, A. (1963) Function of aggregated reticulocyte ribosomes in protein synthesis. Journal of molecular biology, 6, 148–157.

21 Ingolia, N.T., Ghaemmaghami, S., Newman, J.R. and Weissman, J.S. (2009) Genome-wide analysis in vivo of translation with nucleotide resolution using ribosome profiling. Science, 324, 218–223.

22 Andreev, D.E., O'Connor, P.B., Loughran, G., Dmitriev, S.E., Baranov, P.V. and Shatsky, I.N. (2017) Insights into the mechanisms of eukaryotic translation gained with ribosome profiling. Nucleic acids research, 45, 513–526.

23 Ingolia, N.T. (2016) Ribosome Footprint Profiling of Translation throughout the Genome. Cell, 165, 22–33.

24 Andreev, D.E., O'Connor, P.B., Zhdanov, A.V., Dmitriev, R.I., Shatsky, I.N., Papkovsky, D.B. and Baranov, P.V. (2015) Oxygen and glucose deprivation induces widespread alterations in mRNA translation within 20 minutes. Genome biology, 16, 90.

25 Johnson, L.F., Levis, R., Abelson, H.T., Green, H. and Penman, S. (1976) Changes in RNA in relation to growth of the fibroblast. IV. Alterations in theproduction and processing of mRNA and rRNA in resting and growing cells. The Journal of cell biology, 71, 933–938.

26 Gandin, V., Sikstrom, K., Alain, T., Morita, M., McLaughlan, S., Larsson, O. and Topisirovic, I. (2014) Polysome fractionation and analysis of mammalian translatomes on a genome-wide scale. Journal of visualized experiments : JoVE.

27 Picelli, S., Faridani, O.R., Bjorklund, A.K., Winberg, G., Sagasser, S. and Sandberg, R. (2014) Full-length RNA-seq from single cells using Smart-seq2. Nature protocols, 9, 171–181.

28 Langmead, B., Trapnell, C., Pop, M. and Salzberg, S.L. (2009) Ultrafast and memory-efficient alignment of short DNA sequences to the human genome. Genome biology, 10, R25.

29 Ramskold, D., Wang, E.T., Burge, C.B. and Sandberg, R. (2009) An abundance of ubiquitously expressed genes revealed by tissue transcriptome sequence data. PLoS computational biology, 5, e1000598.

30 Robinson, M.D. and Oshlack, A. (2010) A scaling normalization method for differential expression analysis of RNA-seq data. Genome biology, 11, R25.

31 Ritchie, M.E., Phipson, B., Wu, D., Hu, Y., Law, C.W., Shi, W. and Smyth, G.K. (2015) limma powers differential expression analyses for RNA-sequencing and microarray studies. Nucleic acids research, 43, e47.

32 Wright, G.W. and Simon, R.M. (2003) A random variance model for detection of differential gene expression in small microarray experiments. Bioinformatics, 19, 2448–2455.

33 Benjamini, Y., and Hochberg, Y. (1995) Controlling the false discovery rate: a practical and powerful approach to multiple testing. Journal of the Royal Statistical Society Series B57, 289–300.

34 Larsson, O., Sonenberg, N. and Nadon, R. (2010) Identification of differential translation in genome wide studies. Proceedings of the National Academy of Sciences of the United States of America, 107, 21487–21492.

35 Falcon, S. and Gentleman, R. (2007) Using GOstats to test gene lists for GO term association. Bioinformatics, 23, 257–258.

36 Gregory R. Warnes, B.B., Lodewijk Bonebakker, Robert Gentleman, Wolfgang Huber Andy Liaw, Thomas Lumley, Martin Maechler, Arni Magnusson, and Steffen Moeller, M.S.a.B.V. (2016).

37 Bjur, E., Larsson, O., Yurchenko, E., Zheng, L., Gandin, V., Topisirovic, I., Li, S., Wagner, C.R., Sonenberg, N. and Piccirillo, C.A. (2013) Distinct translational control in CD4+ T cell subsets. PLoS Genet, 9, e1003494.

38 Mao, Y.van Hoef, V., Zhang, X., Wennerberg, E., Lorent, J., Witt, K., Masvidal, L., Liang, S., Murray, S., Larsson, O, et al. (2016) IL-15 activates mTOR and primes stress-activated gene expression leading to prolonged antitumor capacity of NK cells. Blood, 128, 1475–1489.

